# Living in the concrete jungle: carnivore spatial ecology in urban parks

**DOI:** 10.1101/2020.06.09.142935

**Authors:** Siria Gámez, Nyeema C. Harris

## Abstract

People and wildlife are living in an increasingly urban world, replete with unprecedented human densities, sprawling built environments, and altered landscapes. Such anthropogenic pressures can affect multiple processes within an ecological community, from spatial patterns to interspecific interactions. We tested two competing hypotheses, human shields versus human competitors, to characterize how humans affect the carnivore community using multi-species occupancy models. From 2017-2020, we conducted the first camera survey of city parks in Detroit, Michigan, and collected spatial occurrence data of the local native carnivore community. Our 12,106-trap night survey captured detected data for coyotes (*Canis latrans*), red foxes (*Vulpes vulpes*), gray foxes (*Urocyon cinereoargenteus*), raccoons (*Procyon lotor*), and striped skunks (*Mephitis mephitis*). Overall occupancy varied across species (Ψ_coyote_=0.40, Ψ _raccoon_=0.54, Ψ_red fox_ =0.19, Ψ_striped skunk_ =0.09). Contrary to expectations, humans did not significantly affect individual occupancy for these urban carnivores. However, co-occurrence between coyote and skunk only increased with human activity. The observed positive spatial association between an apex and subordinate pair supports the human shield hypothesis. Our findings demonstrate how urban carnivores can exploit spatial refugia and coexist with humans in the cityscape.

## INTRODUCTION

Cities are highly heterogeneous landscapes of risk and reward, borne of unique interactions between anthropogenic and ecological processes (Alberti et al., 2003; Liu et al., 2007). As urbanization and land cover conversion rates continue to increase worldwide, cities have emerged as a new and unique habitat for wildlife. By 2050, over half of the global human population will live in a city while urban development is projected to grow by 120 million hectares globally by 2030 (Mcdonald et al., 2018; United Nations, 2018). Cities can be a source or a sink for mammal species, a duality driven by both increases in availability of food sources and risks of mortality (Bateman, & Fleming, 2012; Lepczyk et al., 2017; Lamb et al., 2020). For example, cougars (*Puma concolor*) in an urban-wildland system in Colorado successfully exploited anthropogenic food sources, yet faced a 6.5% increase in mortality risk in developed areas (Moss et al., 2016). Wildlife responses to the built environment are unsurprisingly driven by humans themselves and their induced modifications to landscapes through food provisioning, artificial habitat and light, and roads (Clucas, & Marzluff, 2011; Riley et al., 2014; Gaston et al., 2017).

Anthropogenic pressures affect wildlife communities from intraguild interactions down to behavioral shifts in individual species. Perturbations to higher trophic levels can have cascading impacts on ecosystem processes, which underscores the need to understand how carnivores respond to human activities (Terborgh, 2010; Ripple et al., 2014b). A study in the city of Chicago found that raccoons (*Procyon lotor*) comprised a larger relative proportion of the mesopredator community in urban compared to rural sites, irrespective of patch size (Prange, & Gehrt, 2004). Individual species responses to human activity are varied and depend on each species’ life history traits and behavioral tolerance of human encounters. Cottontail rabbits (*Sylvilagus floridanus*) in an urban area were more vigilant at sites where coyotes were absent, suggesting humans are a third “player” in a predator-prey-human system (Gallo et al., 2019). Evidently, urban systems produce a suite of complex and synergistic changes to ecological communities.

Despite evidence that human activity induces complex responses in urban wildlife, there is a dearth of studies that quantify these effects, particularly for terrestrial carnivores. A meta-analysis of urban ecology studies found that only 10.2% of 244 studies quantified large mammal responses to urbanization and only 6% of all urbanization metrics employed in these studies explicitly considered humans (Moll et al., 2019). Worldwide population declines and range contraction in carnivores highlight the urgency to assess how spaces dominated by humans alter interactions within ecological communities (Ceballos, & Ehrlich, 2002; Ripple et al., 2014a).

We leveraged a North American carnivore guild comprised of coyotes, raccoons, red foxes, gray foxes (*Urocyon cinereoargenteus)*, and striped skunks to investigate how human activity influences spatial ecology within the community. We implemented the first camera survey of city parks in Detroit, Michigan from 2017-2020 to study the city’s carnivores. By directly measuring human activity and not proxies of human pressure such as housing density, we explicitly disentangled the effects of humans on wildlife from those related to the built environment.

In characterizing human effects many relationships may occur on carnivore communities. We underscore two pervasive theoretical frameworks in the literature with distinct expectations for mammalian community response to fine-scale human activity, but recognize these hypotheses are not mutually exclusive (Figure 1). The human shield hypothesis (HSH) argues that humans differentially exert top-down pressure on the apex (hereafter dominant) predator in the system, indirectly benefitting the subordinate competitors and facilitating greater spatial overlap between humans and subordinate species (Shannon et al., 2014; Moll et al., 2018). Anthropogenic pressure is known to mediate intraguild interactions (Berger, 2007; Muhly et al., 2011; Gallo et al., 2019). For instance, red foxes (*Vulpes vulpes*) have been shown to exploit highly developed core urban areas as spatial refugia to avoid their dominant coyote (*Canis latrans*) competitors (Moll et al., 2018). A contrasting approach frames humans as competitors (HCH), asserting that anthropogenic pressure affects multiple species irrespective of their trophic level or dominance hierarchy (Chapron, & López-Bao, 2016; Farris et al., 2017). According to HCH, human presence is functionally similar to antagonism from another competitor in the guild, resulting in increased vigilance, competitive exclusion, and spatial avoidance across the entire community (Clinchy et al., 2016; Gallo et al., 2019; Suraci et al., 2019).

**Figure 1.**
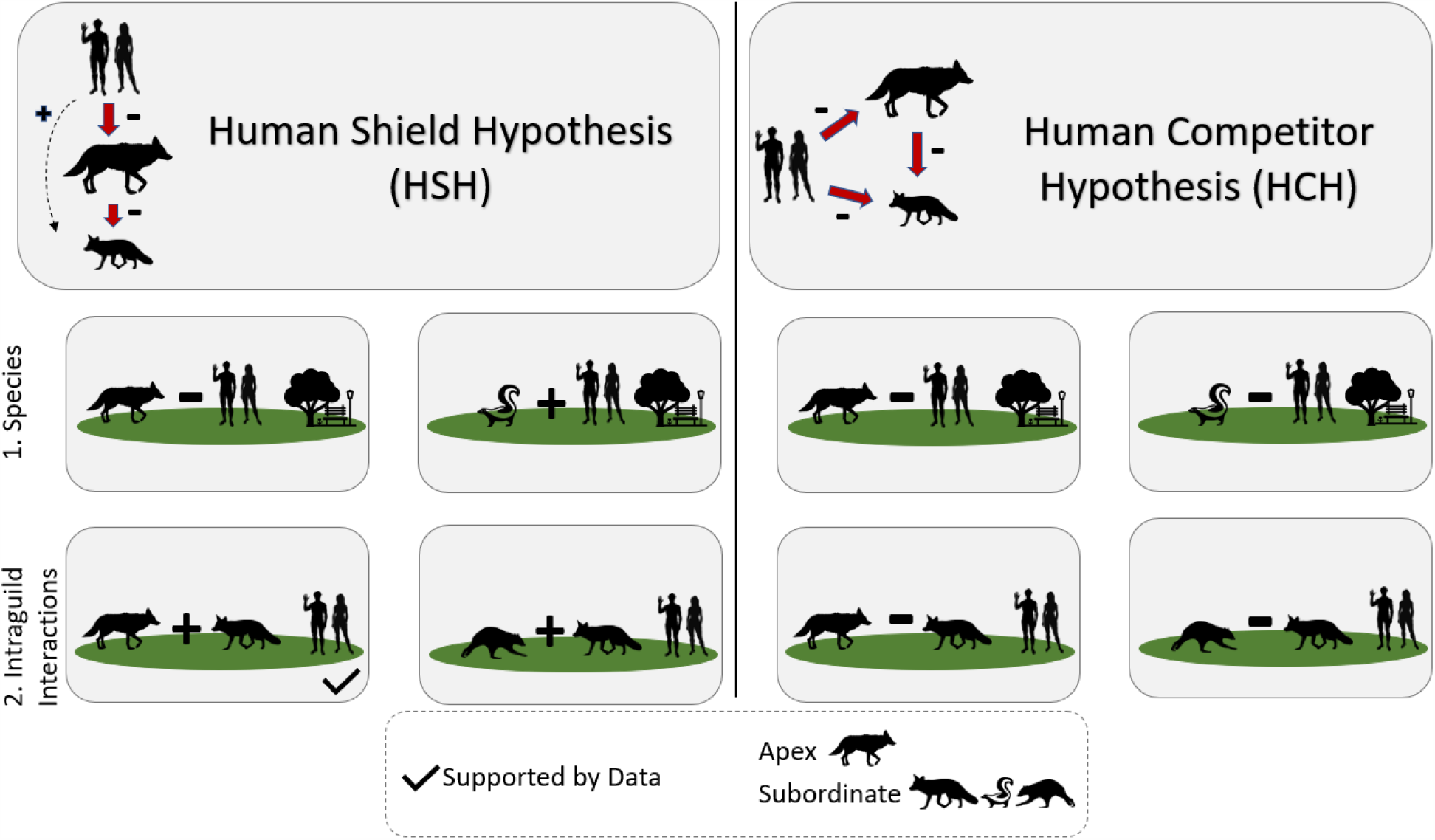
Conceptual framework for the effects of humans on (1) individual carnivore species and (2) pairwise intraguild interactions under two hypotheses: humans as shields (HSH) and humans as competitors (HCH). HSH1: Human reduce dominant carnivore occupancy and increase subordinate carnivore occupancy. HSH2: Increased conditional occupancy for dominant-subordinate and subordinate-subordinate species pairs. HCH1: Humans reduce occupancy for both dominant and subordinate carnivore species. HCH2: Reduced conditional occupancy for dominant-subordinate and subordinate-subordinate species pairs.

Here, we addressed the following questions to test whether effects from anthropogenic pressures on urban carnivores align with expectations of the HSH or HCH: 1) how do humans influence space use of individual carnivore species? 2) how are pairwise interactions affected by human activity within a competing carnivore guild?

With HSH, subordinate mesopredators will exploit the spatial refugia created by human top-down pressures on dominant predators and spatially overlap with humans at the park scale (Geffroy et al., 2015; Moll et al., 2018). We categorized raccoon, red fox, and skunk as subordinate based on body size, trophic level, and known antagonistic interaction with coyotes, which we categorized as dominant (Fedriani et al., 2000). First, for individual species we expected human activity to reduce dominant carnivore occupancy and concurrently increase subordinate carnivore occupancy (Berger, 2007; Muhly et al., 2011). Second, for pairwise interactions we expected human activity to increase dominant-subordinate and subordinate-subordinate carnivore species pair conditional occupancy (Smith et al., 2018).

Conversely, with HCH humans can function like a superior competitor, reduce the niche space and thus spatially displace carnivores regardless of whether they are dominant or subordinate (Everatt et al., 2019). First, for individual species we expected human activity to reduce both dominant and subordinate carnivore occupancy (George, & Crooks, 2006). Second, for pairwise interactions we expect human activity to reduce both subordinate-dominant and subordinate-subordinate carnivore conditional occupancy (Magle et al., 2014).

## MATERIALS & METHODS

### Study Area

We surveyed the carnivore community at 24 urban parks throughout Detroit, a ∼ 370 km^2^ city in southeastern Michigan, USA using remotely triggered cameras (Figure 2). Parks sampled as part of our study represent 51% of the total area of the green space in city parks (City of Detroit, 2015). The Detroit River runs along the city’s southern boundary and the Rouge River runs through the southwestern districts including an automotive manufacturing plant. Over the last 70 years, Detroit has experienced a substantial population decline, dropping from 1.8 million residents at the height of its industrial era in 1950 to approximately 673,000 residents in 2016 (U.S. Census Bureau, 2016). Current human population density is 1,819/km compared to 2,374/ km found in the smaller mid-western city of Milwaukee, Wisconsin. The number of empty lots peaked at roughly 120,000 vacant lots (33% of total parcels) and 48,000 abandoned buildings (13% of parcels) in 2010 (Detroit Residential Parcel Survey, 2010; Raleigh, & Galster, 2015). Over time, many empty lots have progressed through early successional stages and have even developed enough vegetative cover to support small mammal populations, a key prey source for urban carnivores (Bateman, & Fleming, 2012).

**Figure 2.**
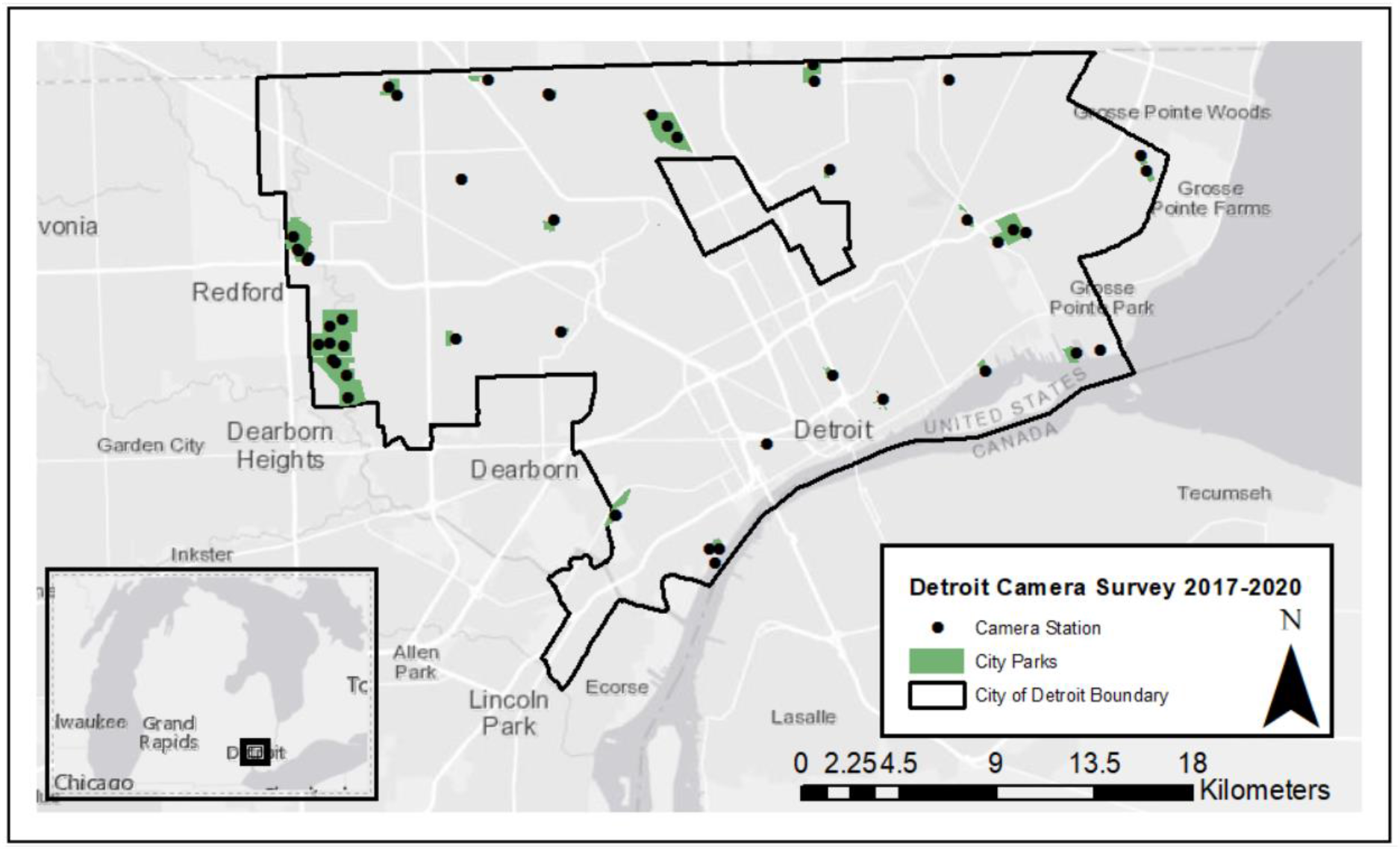
Study area – City of Detroit, Michigan. Shaded green polygons represent the 24 city parks in the Detroit Metro Parks system included in the analysis while black dots denote camera stations in the study.

### Camera Survey

We conducted a 3-year, non-invasive survey by installing motion-triggered trail cameras (Reconyx© PC 850, 850C, 900, 900C) throughout city parks during the fall-winter season in Detroit (November 2017-March 2018, November 2018-February 2019, November 2019 – March 2020). We deployed 39 stations across 23 city parks in 2017, 41 stations across 24 parks in 2018, and 36 stations at 23 parks in 2019. The same parks were sampled across 2017-2020, with the exception of 2018 where a new site was added. We selected parks to ensure representation of ecological and anthropogenic features such as park size, vegetation cover, distance to water, trails, and built infrastructure such as visitor pavilions and playgrounds. Site selection for camera placement within parks was based on animal sign such as tracks, scat, or natural trails to improve detection of carnivores and their prey. Unbaited cameras were placed approximately 0.5 m from the ground on trees >10 cm in diameter, following standard protocol for mesocarnivore camera trap studies (Cove et al., 2012). Camera settings were set to high sensitivity, with three images captured per trigger at 1s intervals, and 15s delay between triggers. For parks with > 1 camera station, the average distance between cameras was 1,416 m, while the average distance between parks was 3,200 m.

After camera retrieval, at least two members of the Applied Wildlife Ecology Lab at the University of Michigan classified images to species and confirmed accuracy. Any unresolved photos were classified as “unknown” and removed from the analysis. We implemented a 30-minute quiet period to account for pseudoreplication and improve independence for analysis, given some animals tend to remain in front of the camera and trigger it multiple times. Domestic dogs and cats were excluded from the analysis as we could not differentiate between feral animals who form part of the local carnivore community and those which were temporarily roaming from their owners. Focal species in this study are relatively common in the eastern United States and comprise a guild that is hierarchically structured, ideal for investigating the effects of human presence on the space use of a carnivore community in urban environments.

### Modeling Carnivore Occupancy

Camera trap data was formatted for occupancy analysis by generating weekly detection histories (i.e. presence ‘1’ or absence ‘0’ of each species at each camera location) using the R package ‘camtrapR’ (Niedballa, 2016). We used a Bayesian multi-species occupancy modeling approach and fit a series of candidate models to test hypotheses about anthropogenic effects on individual carnivore species and intraguild interactions (Rota et al., 2016). The latent occupancy state (Ψ) was modeled as a multivariate Bernoulli random variable (MVB), where Z_i_ = {Z_i1_, Z_i2_, Z_i3_, Z_i4_} represents the 4-dimensional vector of binary detection data for the four focal carnivores (Dai et al., 2013). Each occupancy state represents possible scenarios, i.e. Ψ_1111_ denotes all four species are present, Ψ_0000_ denotes all carnivores are absent.

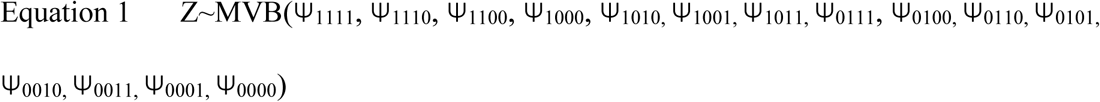

Interaction parameters for were included in the model to calculate individual occupancy estimates as well as conditional probabilities for each species. Conditional probabilities reflect occupancy estimates given the presence of a competitor (i.e. occupancy estimate of red fox, given the presence of coyote). We modeled natural parameters (f_1_, f_2_, f_3_, f_4_, f_12_, f_13_, f_14_, f_23_, f_24_, f_34_) as linear functions to obtain the probability of each community state, where each subscript denotes one of the four species: 1=coyote, 2=raccoon, 3=red fox, 4=skunk, following derivations of (Rota et al., 2016). The number of grey fox detections was insufficient for inclusion in the occupancy models and were excluded from the analysis (n=9 detections at 1 site) (Mackenzie et al., 2002).

The detection process was modeled as a function of covariates including park area (km^2^) (AREA), number of trap nights (TN), camera type (CAM), and NDVI as a measure of vegetation density (VEG); occupancy was modeled as a function of human detections per trap night (HUM), distance to school (km) (DSCH), year (YR), AREA, and NDVI. Top models were selected with an information theoretic approach using Akaike’s Information Criterion (AIC) to identify the model with the lowest ΔAIC and greatest weight (*w*) (Mackenzie, & Bailey, 2004). Models were run using the *unmarked* package in Program R; model fit was assessed using residual sum of squares (RSS) with the parametric bootstrap function ‘parboot’ (Fiske, & Chandler, 2011; R Core Team, 2017). We assessed the effects of human activity on carnivore occupancy using the HUM covariate coefficient estimate (β_HUM_) from the model with the lowest ΔAIC. A negative β_HUM_ signifies that humans decreased individual species or conditional pairwise occupancy, while a positive β_HUM_ indicated that human activity increased these occupancy estimates. We calculated the 95% confidence interval for each β_HUM_ to determine whether it was a strong predictor of occupancy and concluded that intervals overlapping zero were poor predictors and thus did not affect occupancy.

## RESULTS

Our 12,106-trap night survey yielded detections of coyotes (n=220), raccoons (n=1,496), red foxes (n=88), grey foxes (n=11), and striped skunks (n=38) in a highly urbanized landscape from 2017-2020. The proportion of sites occupied (uncorrected for imperfect detection) varied by species (Table 1). We recorded 1,103 human detections at 25 parks, resulting in a naïve occupancy estimate of 0.64 based on the proportion of occasions where humans were present across our study period.

**Table 1.**
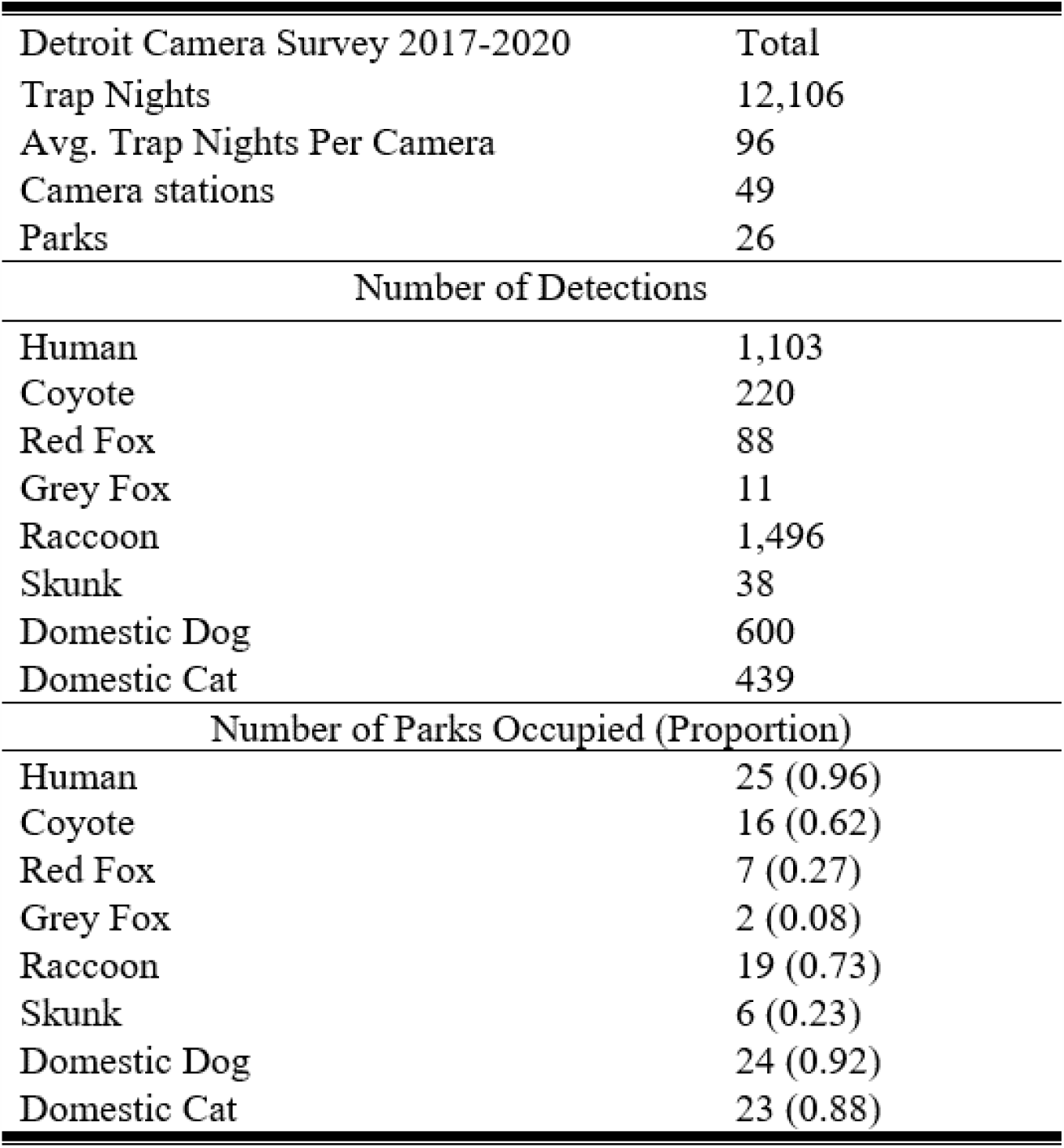
Summary of 2017-2020 DMP camera survey including total and average number of trap nights, camera stations, parks, detections for all species, as well as number and proportion of parks occupied by each species (i.e., naïve occupancy).

### Species Response to Human Activity

Contrary to expectations of both HCH and HSH, human activity did not significantly alter single species occupancy estimates for any of the four focal carnivore species (Figure 3A). Human β_HUM_ coefficients derived from the top performing model for individual species occupancy were: coyote (β_HUM_=-0.13, 95% CI:-11.6, 11.3), raccoon (β_HUM_=-1.3, CI:-3.9, 1.3), red fox (β_HUM_=1.5, CI:-1.5, 4.5), and skunk β_HUM_=-2.9, CI:-14.8, 8.9). Other environmental variables were strong predictors of individual species occupancy. For example, DSCH increased occupancy for raccoons (β_DSCH_=0.001, CI:0.0008, 0.0012). VEG and AREA were negative predictors for red foxes, while they were positive and negative predictors for coyotes, respectively (Table 2)

**Table 2.** Summary of candidate multispecies occupancy models with covariates including year (YR), park area (km^2^) (SIZE), human detections per trap night (HUM), distance to school (km) (DSCH), and NDVI (VEG) as a measure of vegetation density. Dot models (.) denote null (i.e. constant) detection (*ρ*) or occupancy(ψ). Akaike’s Information Criterion (AIC), delta AIC, AIC model weight (*w*), and model goodness of fit residual sum of squares (RSS *p-value*) are listed.

| Model | Equation | K | AIC | ΔAIC | $w$ | RSS $p$-value |
| --- | --- | --- | --- | --- | --- | --- |
| 3 | ρCoy (VEG); ΨCoy (AREA+VEG+HUM),<br>ρRac (.); ΨRac (DSCH+HUM),<br>ρFox (VEG); ΨFox (AREA+VEG+HUM),<br>ρSku (.); ΨSku (HUM),<br>ρCoy-Rac (.); ΨCoy-Rac (AREA+HUM),<br>ρCoy-Fox (.); ΨCoy-Fox (AREA+VEG+HUM),<br>ρCoy-Sku (.); ΨCoy-Sku (AREA+HUM),<br>ρRac-Fox (.); ΨRac-Fox (VEG+HUM),<br>ρRac-Sku (.); ΨRac-Sku (HUM),<br>ρFox-Sku (.); ΨFox-Sku (HUM) | 36 | 3356.85 | 0.00 | 0.74 | < 0.01 |
| 1 | ρCoy (.); ΨCoy (AREA+VEG),<br>ρRac (.); ΨRac (DSCH+HUM),<br>ρFox (.); ΨFox (AREA+VEG),<br>ρSku (.); ΨSku (.),<br>ρCoy-Rac (.); ΨCoy-Rac (AREA+HUM),<br>ρCoy-Fox (.); ΨCoy-Fox (AREA+VEG),<br>ρCoy-Sku (.); ΨCoy-Sku (AREA),<br>ρRac-Fox (.); ΨRac-Fox (VEG+HUM),<br>ρRac-Sku (.); ΨRac-Sku (DSCH),<br>ρFox-Sku (.); ΨFox-Sku (DSCH) | 29 | 3359.52 | 2.67 | 0.19 | < 0.01 |
| 2 | ρCoy (NDVI); ΨCoy (AREA+VEG),<br>ρRac (.); ΨRac (DSCH),<br>ρFox (NDVI); ΨFox (AREA+VEG),<br>ρSku (.); ΨSku (.),<br>ρCoy-Rac(.); ΨCoy-Rac (AREA),<br>ρCoy-Fox(.); ΨCoy-Fox (AREA+VEG),<br>ρCoy-Sku(.); ΨCoy-Sku (AREA),<br>ρRac-Fox(.); ΨRac-Fox (VEG),<br>ρRac-Sku(.); ΨRac-Sku (.),<br>ρFox-Sku(.); ΨFox-Sku (.) | 26 | 3363.63 | 6.78 | 0.026 | < 0.01 |

**Figure 3.**
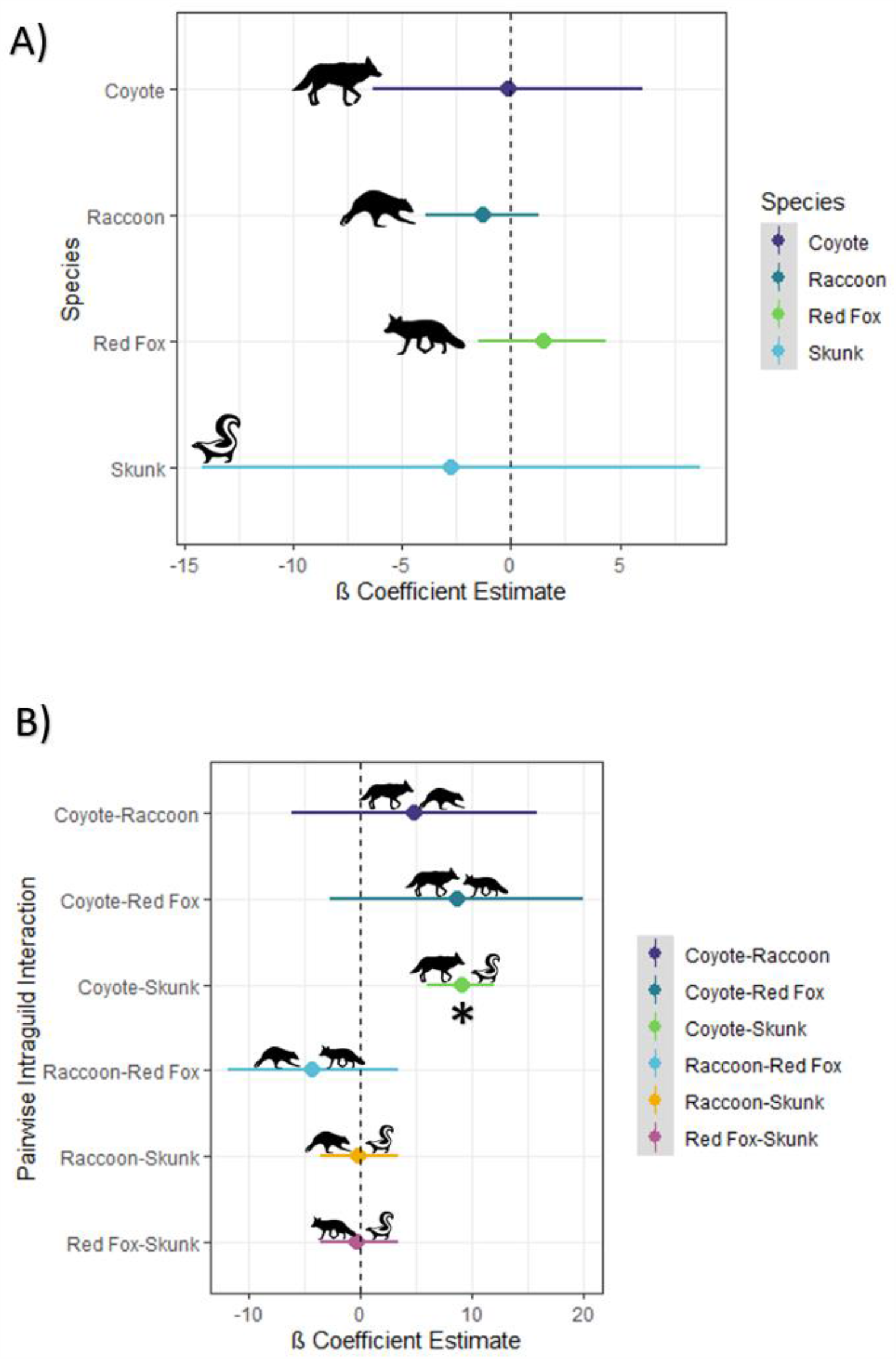
Human effects on A) Individual carnivore species and B) Pairwise intraguild interactions using β coefficient estimates for the human trap night (HUM) covariate and 95% confidence intervals shown from top model. Asterisk denotes significant effect with interval not overlapping 0.

### Pairwise Intraguild Interaction Response to Human Activity

Human activity altered how urban carnivores use space with respect to their intraguild competitors. The inclusion of HUM as a covariate for all individual and pairwise interaction occupancy estimates greatly improved model inference, demonstrating fine-scale measures of human activity are informative in explaining carnivore occupancy. Human coefficient estimates for coyote-raccoon (β_HUM_=4.8, CI:-6.2, 15.8), coyote-red fox(β_HUM_=8.7, CI:-2.6, 20.0), raccoon-red fox (β_HUM_=-3.9, CI:-11.3, 3.5), raccoon-skunk (β_HUM_=-0.12, CI:-0.4, 0.2), red fox-skunk (β_HUM_=-0.2, CI:-1.0, 0.6), overlapped zero, signaling that humans were weak predictors of conditional occupancy for these species pairs. However, human activity significantly increased the likelihood that skunks would occupy an area where coyotes were present (β_HUM_=9.0, CI:8.7, 9.2) in support of the human shield hypothesis with intraguild interactions between dominant and subordinate species pair (HSH, Figure 3B).

## DISCUSSION

Our study provides necessary insights into how human activity affects space use in the carnivore community in Detroit parks. Common proxies for human activity derived from landscape-level metrics of urbanization such as housing density, percent impervious surfaces, or road density may not capture the resolution necessary to determine fine-scale consequences for wildlife (Tablado, & Jenni, 2017). The inclusion of indices of direct human activity is therefore needed in future urban ecology studies to disentangle the effects of humans from the built environment (Nickel et al., 2020). Further, the design of parks and urban green spaces should include considerations of how human presence shapes animal communities to adequately balance the needs of people and wildlife.

Contrary to our expectations, humans did not influence single species occupancy, a result which neither supports HSH or HCH. Our hypothetical framework hinged on either a positive or negative effect of humans on urban carnivore space use. However, our observed lack of effect for individual species may reflect an alternate hypothesis wherein humans do not directly influence space use at a fine scale, likely as a result of heightened behavioral plasticity in urban adapters (Lowry et al., 2013). Key resources such as den sites and prey availability may be intrinsically concentrated in urban green spaces, given that they are essentially habitat fragments embedded in an urban matrix, thus funneling carnivores into parks regardless of human activity (Marzluff, 2005). Our results are consistent with a growing body of evidence that underscores the importance of urban greenspaces to ensure long-term persistence carnivore community (Gallo et al., 2017).

We found an influence of human activity altering intraguild interactions for only one species pair. Spatial interaction between coyotes and skunks increased significantly with human activity, indicating that humans effectively shield skunks from coyotes consistent with HSH. As the smallest carnivore in our study, they face antagonism from coyotes ranging from interference competition to direct killing (Fisher, & Stankowich, 2018). To date, there has been little evidence of skunks spatially avoiding coyotes, thus we present a novel example of how skunk spatial ecology is mediated by human activity (Prange, & Gehrt, 2007; Ritchie, & Johnson, 2009). The long-term effects of such increased spatial overlap are unknown between this species pair. If humans continue to shield skunks from dominant competitors, the result is a net positive for skunks as coexistence is facilitated. Alternatively, if the interaction strength between humans and coyotes weakens, yet skunks continue to use humans as a shield, they could potentially be lured into more frequent antagonistic interactions with coyotes and face increased mortality risk, akin to an ecological trap (Robertson, & Hutto, 2006; Bateman, & Fleming, 2012). Carnivores may leverage the temporal niche dimension to avoid competitive exclusion, though our study did not explore this aspect of urban carnivore ecology and represents a fruitful future direction.

If humans function as shields and facilitate spatial overlap in the carnivore community, then urban areas could serve as key refugia in cities and increase co-existence in an otherwise highly competitive guild. Paradoxically, the human shield hypothesis indicates that urbanization does not inherently result in biodiversity loss at the patch scale, given that subordinate species can exploit refugia (Moll et al., 2018; Lewis et al., 2019). However, biodiversity loss due to urbanization at both the landscape and global scale remains a concern for conservation efforts (Mcdonald et al., 2013; Mcintyre, 2014; Lewis et al., 2015).

In addition to wild carnivores, domestic dogs and cats were commonly detected and often without human company. Human affiliates such as dogs and cats are widely recognized as having significant detrimental effects on wildlife (Lenth et al., 2008; Vanak, & Gompper, 2009; Loss et al., 2013). Despite this, the distinction between free roaming, feral, or simply temporarily off leash remains unresolved in our study system. As a result, we were unable to determine whether these human affiliates were true long-term members of the local carnivore assembly and thus were excluded from the analysis. Leashed dogs sometimes harass and injure wildlife, including some of the focal species in this study (Hughes, & Macdonald, 2013). However, how these antagonistic interactions determine the composition, structure, and distribution of carnivore communities in urban spaces is not well understood. Because dogs are one of the most widely distributed terrestrial carnivores, filling this knowledge gap should be a key consideration for future studies to better inform natural resource managers seeking to mitigate their effects on wildlife (Gehrt et al., 2010).

Our study provides insight into the intraguild interactions of an urban carnivore community in a mid-sized city whose population has declined over the last 70 years. Homogenization patterns expected in urban systems do not necessarily scale down to smaller cities (Collins et al., 2002). Moreover, the historical trajectory and relationship between population decline, housing vacancy, and vegetation varies by city (Schwarz et al., 2018).

Notably, this emigration of people from the city of Detroit is complex and tied to various historical socioeconomic biases (Xie et al., 2018). Further, we recognize that how people are distributed in the city and who has access to green spaces is not equitable and a consequence of discriminatory housing and city planning policies that impact a myriad of ecological processes (Watkins, & Gerrish, 2018; Schell et al., 2020). This economic and racial segregation of neighborhoods introduces a bias to our understanding human-wildlife interactions in cities (Alberti et al., 2020). The location, maintenance level, and surrounding characteristics of parks are also unequally distributed in cities, this inequity can potentially attract or deter carnivore species and confound the interpretation of our results (Elliott et al., 2019; Huang et al., 2020). Further studies are needed to disentangle urban carnivore coexistence patterns and resource availability from socioeconomic factors driven by racial disparities and environmental injustices (Wilson et al., 2008).

Finally, our study could inform how natural resource managers and city planners approach urban design and offer opportunities for collaboration with ecologists. Given that urban carnivores seek spatial refuge from human activity hotspots, future park designs could incorporate wildlife zones where the use of walking trails diverts human foot traffic around rather than through important habitat (Hess et al., 2014). Urban planners are thus tasked with promoting access to natural areas for the public, while still conserving habitat for wildlife. City parks are an important resource for urbanites and provide recreational, cultural, psychological and physiological benefits to visitors (Soga, & Gaston, 2020). Therefore, finding a balance between the wellbeing of people and wildlife is a fundamental challenge of the 21^st^ century (Chawla, 2015; Rigolon, 2016; Liu et al., 2017).

## ACKNOWLEDGEMENTS

First, we recognize implementing our field research with camera traps was conducted on lands originally belonging to the People of the Three Fires. Our work is not human subjects research requiring IRB review, though we remain grateful to authorities granting permission for our research and their efforts to manage coupled human-natural ecosystems. Our sincere thanks to members past and present of the Applied Wildlife Ecology (AWE) Lab at the University of Michigan, specifically K. Mills, R. Malhotra, S. Lima, S. Bower, and G. Gadsden who contributed to the data collection, image sorting, ArcGIS expertise, and logistical support of this project. We thank our partners at the Detroit Zoological Society for their financial support and the City of Detroit for collaboration, permits, and access to the parks in our study. We thank all of our volunteers for their assistance with camera checks as well as the *Michigan ZoomIN* online community for their contribution to image classification.

